# SARS-CoV-2 PLpro whole human proteome cleavage prediction and enrichment/depletion analysis

**DOI:** 10.1101/2021.10.04.462902

**Authors:** Lucas Prescott

## Abstract

A novel coronavirus (SARS-CoV-2) has caused a pandemic that has killed millions of people, worldwide vaccination and herd immunity are still far away, and few therapeutics are approved by regulatory agencies for widespread use. The coronavirus 3-chymotrypsin-like protease (3CLpro) is a commonly investigated target in COVID-19, however less work has been directed toward the equally important papain-like protease (PLpro). PLpro is less characterized due to its fewer and more diverse cleavages in coronavirus proteomes and the assumption that it mainly modulates host pathways with its deubiquitinating activity. Here, I extend my previous work on 3CLpro human cleavage prediction and enrichment/depletion analysis to PLpro.[1] Using three sets of neural networks trained on different taxonomic ranks of dataset with a maximum of 463 different putative PLpro cleavages, Matthews correlation coefficients of 0.900, 0.948, and 0.966 were achieved for *Coronaviridae*, *Betacoronavirus*, and *Sarbecovirus*, respectively. I predict that more than 1,000 human proteins may be cleaved by PLpro depending on diversity of the training dataset and that many of these proteins are distinct from those previously predicted to be cleaved by 3CLpro. PLpro cleavages are similarly nonrandomly distributed and result in many annotations shared with 3CLpro cleavages including ubiquitination, poly(A) tail and 5’ cap RNA binding proteins, helicases, and endogenous viral proteins. Combining PLpro with 3CLpro cleavage predictions, additional novel enrichment analysis was performed on known substrates of cleaved E3 ubiquitin ligases with results indicating that many pathways including viral RNA sensing are affected indirectly by E3 ligase cleavage independent of traditional PLpro deubiquitinating activity. As with 3CLpro, PLpro whole proteome cleavage prediction revealed many novel potential therapeutic targets against coronaviruses, although experimental verification is similarly required.

## Introduction

Coronavirus genomes contain multiple open reading frames, the largest of which contains two nonstructural cysteine proteases, nsp3/PLpro and nsp5/3CLpro, which cleave its polyprotein into functional proteins. 3CLpro cleaves 11 conserved sights in all coronavirus proteomes, and PLpro cleaves only 2 or 3 sites depending on whether nsp1 is functional. In addition to many other multifunctional domains, PLpro contains two papain-like domains with different properties, the first of which is sometimes inactive.[2]

Most experimental work on PLpro focuses on its deubiquitinating and deISGylating activity[3] and faster kinetics relative to the human homologs USP7 and USP14.[4] Proteins and pathways disrupted by this activity include retinoic acid-inducible gene I (RIG-I), melanoma differentiation-associated protein 5 (MDA5), TNFR-associated factors (TRAF3/6), stimulator of interferon genes (STING), TANK-binding kinase 1 (TBK1), nuclear factor κ-light-chain-enhancer of activated B cells (NF-κB), interferon regulatory factor 3 (IRF3), type-I interferon production, Janus kinases and signal transducers and activators of transcription (JAK-STAT), and collagen expression via transforming growth factor (TGF-β1).[5–16] Separately, simple sequence alignments of the three SARS-CoV-2 cleavages with human proteins ranked by percent identity led Reynolds et al. to propose a few dozen human proteins to be directly cleaved by PLpro.[17] This method does not account for position-dependent residue importance or any physiochemical trends and therefore likely resulted in many false positives, yet they were able to demonstrate *in vitro* cleavage of protein S, MHC-α/β isoforms (MYH6/7), forkhead box P3 (FOXP3), and epidermal growth factor receptor (ERBB4). IRF3,[18] Unc-51 like autophagy activating kinase (ULK1),[19] proto-oncogene SRC, Kelch domain-containing protein 10 (KLHDC10), and bone marrow stromal antigen 1 (BST1)[20] were also independently demonstrated to be cleaved by PLpro *in vitro*.

The high number of known 3CLpro cleavages has allowed for sequence logos and machine learning techniques to be applied to coronavirus and human sequences.[1, 21] PLpro, however, is less characterized because (1) fewer cleavage sites are known, (2) these cleavages are not as conserved as 3CLpro cleavages, and it is unknown whether (3) any alpha- or betacoronaviruses are missing a cleavage before nsp2, (4) all gamma- and deltacoronaviruses are missing a cleavage before nsp2, or (5) whether the many uncharacterized coronaviruses have one or two functional papain-like domains and what differences they may have.

## Methods

### Data Set Preparation

A complete, manually reviewed human proteome containing 20,350 sequences (not including alternative isoforms) was retrieved from UniProt/Swiss-Prot (proteome:up000005640 AND reviewed:yes).[22]

Additional coronavirus polyprotein cleavages were collected from GenBank.[23] Searching for all combinations of “orf/pp” and “1/1a/1ab” within the family *Coronaviridae* returned 12,770 different, complete polyproteins with 463 different cleavages manually discovered using the Clustal Omega multiple sequence alignment server.[24–26] Unlike 3CLpro, no PLpro domain is found in the monotypic *Microhyla letovirus 1* in *Coronaviridae*. Similar domains are characterized in *Arteriviridae* and in some of *Tobaniviridae*, although none of these sequences were aligned here.[27] All unbalanced positive cleavages were used for subsequent classifier training in addition to all other 33,185 uncleaved coronavirus sequence windows with at least two glycines or alanines in the P2, P1, or P1’ positions, totaling 33,648 samples.

### Cleavage Prediction

As in my previous work on 3CLpro,[1] sequence logo-based logistic regression and naïve Bayes classification and physiochemical and one-hot encoded neural networks (NNs) were used for prediction of cleavage sites.[28]

### Enrichment Analysis

Protein annotation, classification, and enrichment analysis was performed using the Database for Annotation, Visualization, and Integrated Discovery (DAVID) 6.8.[29, 30] Tissue (UP_TISSUE and UNIGENE_EST_QUARTILE), InterPro, direct Gene Ontology (GO includes cellular compartment (CC), biological process (BP), and molecular function (MF)), Reactome pathways, sequence features, and keywords annotations were all explored, and only annotations with Benjamini-Hochberg-corrected p-values less than 0.05 were considered statistically significant. Cleaved E3 ubiquitin ligases were matched to their respective substrates for additional enrichment analysis using the UbiNet 2.0 database.[31] All training data, prediction methods, and results can be found on GitHub (https://github.com/Luke8472NN/NetProtease).

## Results

Due to the greater difficulty aligning putative cleavages and more variable domains of nsp3, the assumption that SARS-CoV-2 PLpro can cleave all aligned sites in divergent coronaviruses is less plausible than for 3CLpro. First, some entries in this dataset containing one, two, or four glycines or alanines may be incorrectly shifted by one or two residues because they are difficult to align with PLpro’s consensus [KL]K[GAR][GA]^[GA]. The P1’ glycine or alanine is also not applicable when cleaving isopeptide bonds for deubiquitination, however lysine’s bound side chain somewhat resembles another small, hydrophobic peptide bond. Glycine is the smallest and most similar amino acid, and alanine is the second smallest, also resembling the hydrophobicity of lysine’s side chain (Figure 1) and potentially similarly protecting the oxyanion of the tetrahedral intermediate from water in the tryptophan-containing oxyanion hole.[6] If this is correct, then ubiquitin’s arginine residue should be in the P3 position and not the P2 position as hinted by many sequences aligned here, particularly nsp1/2 cleavages (Figure 2). Molecular docking or crystallography are, however, required to prove these chemical similarities relevant. Additionally complicating the problem, dimensionality reduction of putative cleavages did not cluster by order as well as for 3CLpro, and nsp1/2 cleavages missing from all gamma- and deltacoronaviruses have a distinct sequence logo (Figure 2).

**Figure 1:**
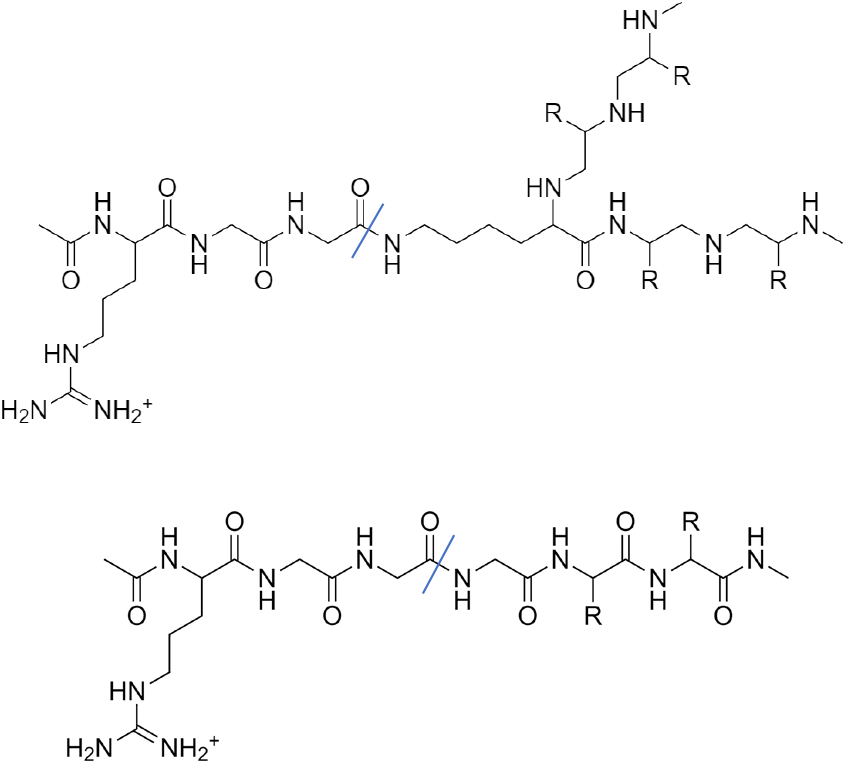
Chemical similarity between (A) ubiquitin-target RGG^K isopeptide bond and (B) RGG^G peptide bond.

**Figure 2:**
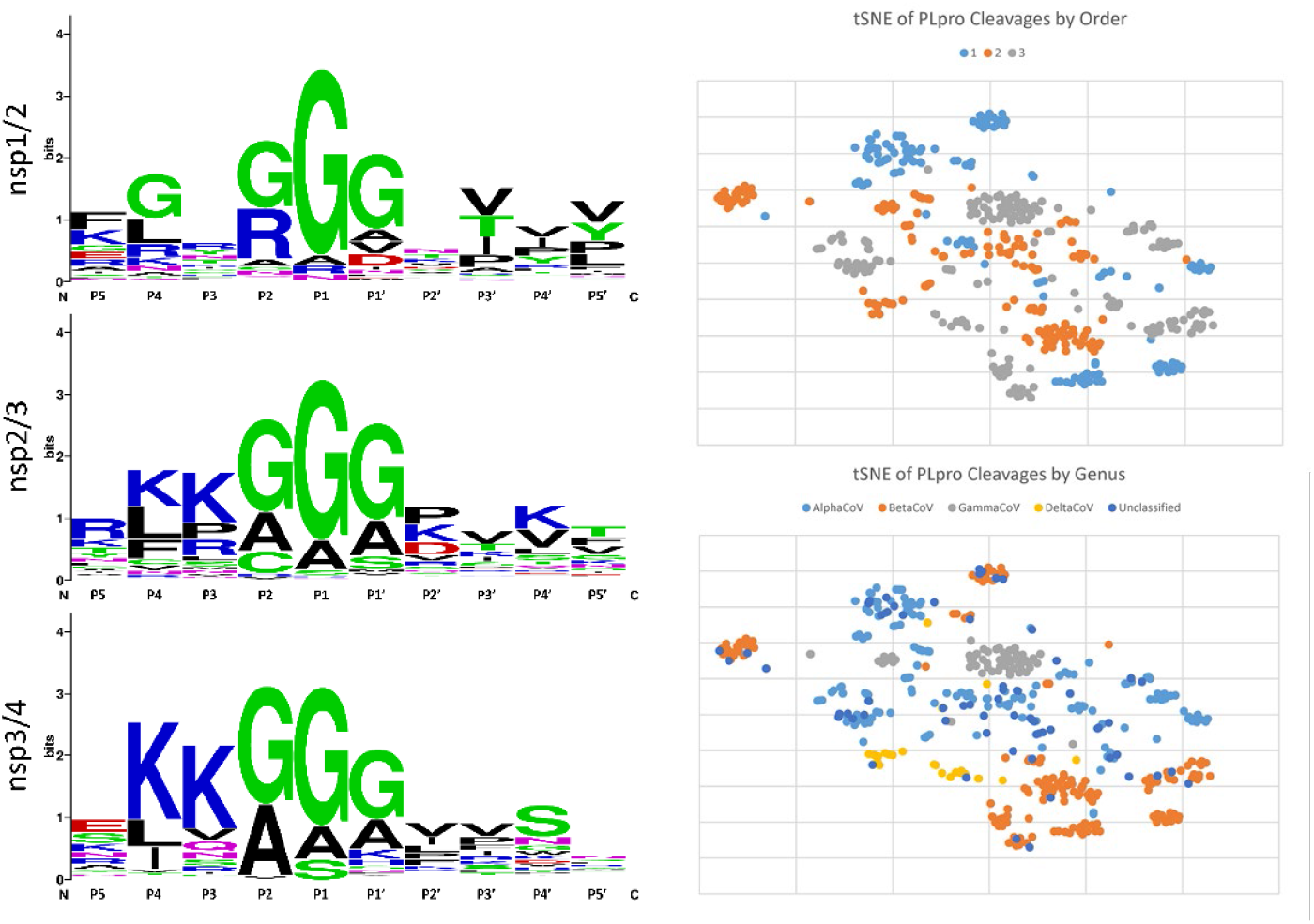
(A) Sequence logos for the three cleavage sites in coronaviruses[32] and (B) one-hot encoded t-SNE colored by order and (C) by genus.[33]

With no easy way to prove functionality or activity of divergent papain-like domains and therefore the suitability of distant cleavages to be included into a single, unweighted dataset, machine learning models were trained on three different taxonomic ranks of data (all *Coronaviridae*, only *Betacoronavirus*, and only *Sarbecovirus*). Predicting sequence-dependent kinetic rates would be even more difficult than this classification given that swine acute diarrhea syndrome coronavirus (SADS-CoV) PLpro catalytic efficiency varies by two orders of magnitude depending of substrate[34] and also given that SARS-CoV-1 and −2 PLpro share 83% identity yet are known to preferentially cleave ubiquitin vs ISG15, respectively.[35] Preferences for isopeptidase over peptidase activity,[36] polyubiquitin chains over ISG15 or monoubiquitin,[37] and K48 over K63 ubiquitin linkages[38] also vary between viral lineages.[39, 40] The distantly related foot-and-mouth disease virus (FMDV) leader protease even has irreversible deISGylating activity by leaving ISG15’s C-terminal diglycine on its target lysine.[41] It is fair to assume, however, that cleavages predicted by all three sets of models can occur in many infections no matter viral lineage. The size and imbalances of these final datasets are listed in Table 1. The greater diversity in the higher taxonomic datasets caused many more random sequences and human proteins to be predicted to be cleaved, however enrichment/depletion analysis proved robust against this.

**Table 1:** Size and imbalance of the three datasets used to train machine learning models and the results of the final NN.

| Rank | *Coronaviridae* | *Betacoronavirus* | *Sarbecovirus* |
| --- | --- | --- | --- |
| nsp1/2 cleavages | 133 | 49 | 16 |
| nsp2/3 cleavages | 153 | 60 | 23 |
| nsp3/4 cleavages | 177 | 46 | 13 |
| Total cleaved | 463 | 155 | 52 |
| Total uncleaved | 33185 | 10013 | 2742 |
| % Cleaved | 1.38% | 1.52% | 1.86% |
| Single network 20% test MCC | 0.895 | 0.934 | 0.988 |
| Ensemble 20% test MCC | 0.900 | 0.948 | 0.966 |
| Random cleavage frequency between 3 Gs or As | 2.00% | 0.54% | 0.07% |
| Random cleavage frequency between $\geq 2$ Gs or As | 0.38% | 0.23% | 0.05% |
| Random cleavage frequency | 0.019% | 0.012% | 0.003% |
| Average residues between random cleavages | 5300 | 8300 | 33000 |
| Human proteins cleaved | 1381 | 980 | 154 |

As with 3CLpro predictions, NNs outperformed all other classifiers no matter taxonomic rank (Figure 3). This performance difference was most noticeable for the whole *Coronaviridae* dataset where single sequence logos cannot learn its multiple populations. Interestingly, physiochemical encoding also proved to describe sequences less efficiently than one-hot encoding and sequence logos for the *Sarbecovirus*-only dataset. The optimized hyperparameters for NNs with one-hot encoding were Adam solver, rectifier (ReLU) activation, 0.000001 regularization, no oversampling, and 1 hidden layer with 100 neurons. Combining networks into ensembles again improved accuracy and stability, so the final results were generated with triplicate 10-fold cross-validated networks with average 20% test Matthews correlation coefficients (MCC) of 0.900, 0.948, and 0.966 for *Coronaviridae*, *Betacoronavirus*, *and Sarbecovirus*, respectively. These accuracies, frequency cleaving random amino acid sequences, and number of human proteins with predicted cleavages are also listed in Table 1.

**Figure 3:**
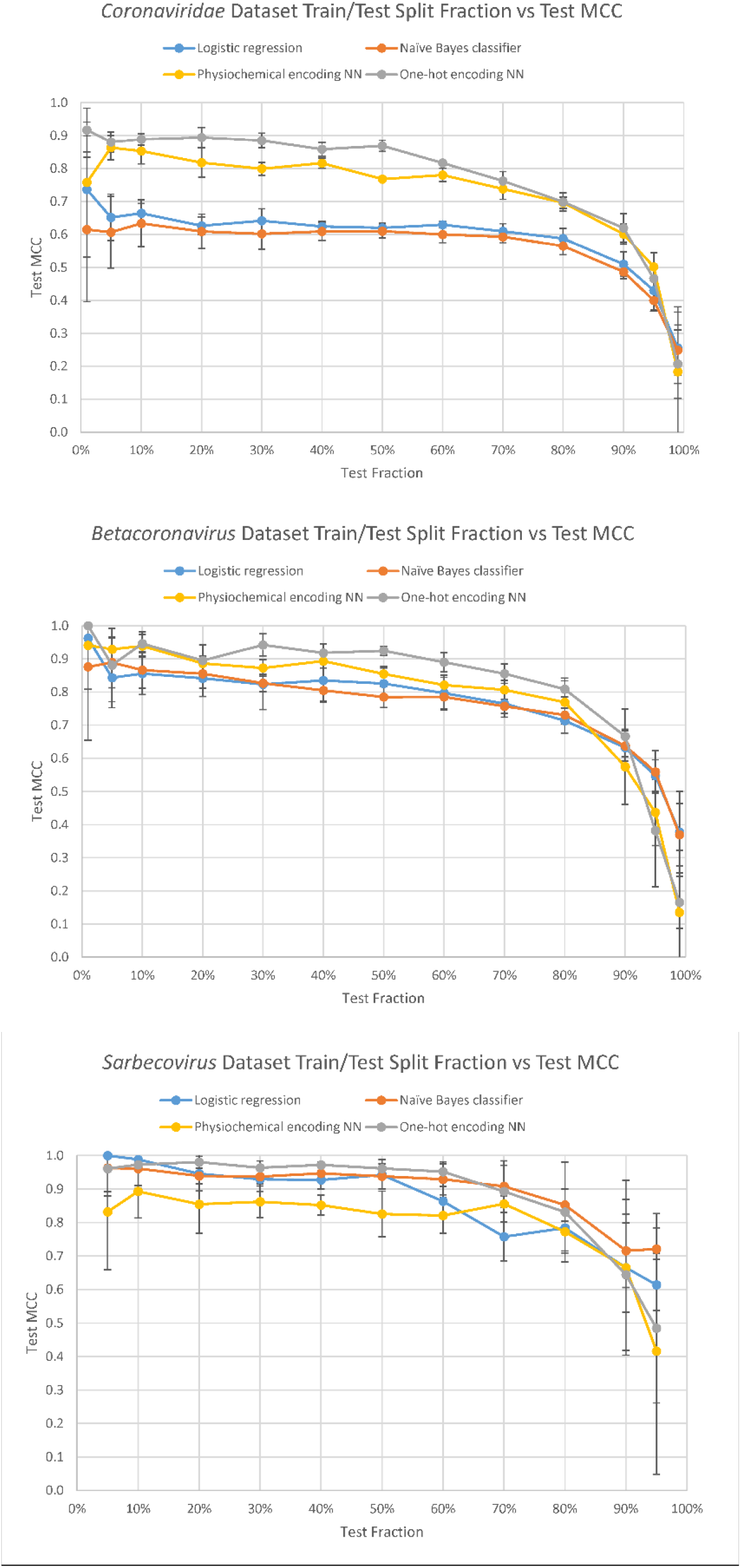
Train/test split fraction vs MCC demonstrating that performance quickly approaches a limit for all classifiers no matter taxonomic rank ((A) *Coronaviridae*, (B) *Betacoronavirus*, (C) *Sarbecovirus)*.

## Discussion

Enrichments and depletions were surprisingly shared between NN replicates, resampled training data, and between the three taxonomic ranks of training data, although the *Sarbecovirus*-specific list returned fewer statistically significant annotations due to its smaller size. As expected from SARS-CoV-2 pathogenesis and from previous 3CLpro results,[1] epithelium (tongue, larynx, pharynx, trachea) and brain were enriched, and olfactory receptors were depleted. As expected from the sequence logo, glycine- and alanine-rich proteins, such as collagens, were also enriched. Cytoskeletal and motor proteins were enriched as with 3CLpro due to their longer lengths, although their cleavage is likely still biologically relevant. In fact, this length bias is even larger in Reynolds et al.’s sequence identity method,[17] and surprisingly few of their predictions were shared in this PLpro prediction dataset, namely protection of telomeres 1 (POT1), membrane-associated guanylate kinase inverted 2 (MAGI2), multiple EGF-like domains 10 (MEGF10), myocardin, inactive RNASE10, amyloid beta A4 precursor protein-binding (APBB2), and myomesin-1. FOXP3 was also the only experimentally validated protein to be successfully predicted here.[18–20] This indicates that, as with 3CLpro prediction, any inaccuracies in prediction methods are amplified when applied to the entire human and must be experimentally verified, however enrichment analyses are robust against this variability.[1]

The ubiquitination pathway was expectedly enriched given one of PLpro’s main functions is to deubiquitinate host proteins. Global effects of PLpro’s preference for particular ubiquitin chains or branchings are poorly characterized, and no peptide bond cleavages within E1, E2, or E3 enzymes are known. Combining these PLpro predictions with my previous 3CLpro dataset,[1] global effects on these pathways become clearer. Of the ten E1 enzymes,[42] two are cleaved by 3CLpro: UBA2 and UBA5. UBA2 is half of the only heterodimer required for SUMOylation and is cleaved in its C-terminal ubiquitin-like domain required for transfer to the only SUMO E2 conjugating protein, UBC9,[43] which is not cleaved by either protease. Interestingly, coronavirus nucleocapsid proteins are intentionally SUMOylated to promote their homo-oligomerization, interfere with cell division, and/or localize to the nucleus or nucleolus to alter viral vs host transcription or ribosomal biogenesis.[44–46] One downstream consequence of disrupted SUMOylation is reduced stability of STING,[47] which is already disrupted by PLpro deubiquitination.[48, 49] TRIM38 (SUMOylating) is not, but SENP2 (deSUMOylating) is cleaved, indicating that STING regulation probably differs between early and late phase infection. PLpro’s additional deubiquitinating activity or cleavage of related enzymes may also change the period of SUMOylation and ubiquitination oscillations, impacting global proteasomal degradation kinetics.[50] Lastly, depletion of E2-conjugated SUMO stores may lead to preferential ubiquitination and degradation of p53[51] by RCHY1 later in infection,[36] which is already directly enhanced by PLpro SARS-unique domain,[52] and IκBα,[53] which activates NF-κB and worsens SARS-related mortality in mice.[54] UBA5, however, is the only UFMylation-specific E1 enzyme with implications in reticulophagy and hematopoiesis, and its only E2 conjugating protein, UFC1, is not cleaved by either protease.[55] First, UFMylation of DDRGK1, a reticulophagy protein activated by ribosome stalling,[56] may play a role in encouraging ribosomal frameshifts by forming double-membrane vesicles (DMVs) utilized by coronaviruses for replication[57] or by compartmentalizing ribosomes with viral RNA inside DMVs as with influenza A virus.[58] Second, like SUMOylation, UFMylation of p53 antagonizes ubiquitination and subsequent degradation.[59] Lastly, UFM1 depletion may encourage macrophage responses to interferon gamma,[60] contributing to the positive feedback with T cells in COVID-19-associated alveolar inflammation.[61] Although none of the few ISG15 enzymes are cleaved by either coronavirus protease, NEDD4, which is likely eventually inhibited by free ISG15 released by PLpro’s deISGylating activity,[62] is predicted to be cleaved by 3CLpro[1] and has been suggested as a therapeutic target for COVID-19 due to its upregulation in infected cells and probable involvement in egress via non-degradative ubiquitination of viral proteins.[63] The counterintuitive proviral activity of NEDD4 yet potential inhibition directly by 3CLpro and indirectly by PLpro hints that its regulation may be infection time dependent.

Interpreting viral intent behind E2 and E3 enzyme cleavages is much more difficult, however similar enrichment analysis can be performed on the substrates specific to cleaved E3 ligases. Combining all predicted cleavages from the three PLpro datasets and the one 3CLpro dataset[1] and applying to UbiNet 2.0 E3-substrate interactions,[31] 38% of substrates were directly cleaved by the viral proteases, and 47% of substrates had E3 ligases cleaved by viral proteases. Because most ubiquitination tags substrates for proteasomal degradation, E3 ligase cleavage would effectively upregulate affected pathways. Although 19% of these cleavages were overlapping and therefore had unknown up- vs downregulation, most noteworthy enrichments were preserved after removal of this overlap from the list of substrates with cleaved E3 ligases. Noteworthy affected pathways include RAP1 related to cell adhesion and junction formation, PI3K/AKT related to cell survival and proliferation, p53 related to apoptosis, clathrin-dependent and -independent endocytosis, Fcγ and Fcɛ receptor signaling, and viral RNA-sensing RIG-I and TLR3 signaling to IRF3/7 and NF-κB and downstream interferon-stimulated STAT1/3. Given these pathways, particularly viral RNA sensing, are affected more directly by many other viral proteins early in infection, pathway activation via cleavage of E3 ligases independent of direct PLpro deubiquitination followed by reduced substrate turnover rate is likely only relevant later in infection once responses to interferon have begun.

In addition to directly deubiquitinating host proteins, PLpro cleavage of lysyl oxidases (LOXL2/3) may disrupt the balance of lysine/acetyllysine/allysine on histones or non-histone proteins[64] to prevent crosslinking or monoubiquitination or disrupt acetylation of ubiquitin itself to prevent polyubiquitination.[65] For example, preventing LOXL3 deacetylation of STAT3 or relatedly cleaving chromodomain-helicase-DNA-binding protein 8 (CHD8), a STAT3 transcriptional repressor, could be part of how SARS-CoV-2 modulates immune cell differentiation[66] or the STAT1/STAT3 ratio.[67]

Outside ubiquitination, RNA binding also stood out as a relevant enrichment because many antiviral pathways attempt to directly degrade viral RNA. Interestingly, the RGG/RG motif present in many RNA-binding proteins[68] is likely enriched in cleavages due to their similarity to PLpro’s sequence logo. Noteworthy cleaved proteins with RNA binding ability include poly(A) tail-associated Ras GTPase-activating protein-binding protein 2 (G3BP2), cleavage and polyadenylation specificity factors (CPSF5/7), zinc finger C3H1 domain-containing protein (ZFC3H1), and zinc finger CCHC domain-containing protein 14 (ZCCHC14) and 5’ cap-associated interferon-induced protein with tetratricopeptide repeats 2 (IFIT2) and dual specificity phosphatase 11 (DUSP11). G3BP2 is involved in stress granule formation and documented to be inhibited by SARS-CoV-2 nucleocapsid protein.[69, 70] Cleavage of RNA cleavage factor Im subunits CPSF5/7 likely contributes to enhanced viral RNA translation by trapping host pre-mRNA in the nucleus as with influenza virus.[71] ZFC3H1 is a subunit of the poly(A) tail exosome targeting (PAXT) complex which targets longer and more polyadenylated RNAs than the traditional nuclear exosome targeting (NEXT) complex[72] with subunit ZCCHC14 also cleaved by PLpro and others cleaved by 3CLpro.[1] In addition to variable length polyadenylation over infection and between whole genomes and subgenomic mRNAs,[73, 74] time-dependent antagonism of these two complexes by PLpro and 3CLpro may be another method coronaviruses control their relative rates of replication vs translation, the ratios of their subgenomic mRNAs, or the ratio of orf1a to orf1ab and resulting proteins. Given hepatitis B virus (HBV) and human cytomegalovirus (HCMV) exploit ZCCHC14 to elongate their poly(A) tails with errors to enhance stability against CCR4-NOT, coronaviruses should also be experimentally surveyed for similar not-so-pure poly(A) tails.[75] In fact, CCR4-NOT subunits 1 and 4, the latter with E3 ubiquitin ligase activity, are cleaved by 3CLpro.[1] In the cytoplasm, this cleavage probably stabilizes viral RNA and viral translation, but disrupting E3 ligase activity in the nucleus would stabilize its target JARID1C, a histone H3 K4 demethylase, globally repressing host transcription so that ribosomes can be directed more toward viral transcripts.[76] As for the two 5’ cap-associated proteins, interferon-stimulated IFIT2 inhibits eukaryotic initiation factor 3 (eIF3)-mediated translation of viral RNAs without cap 2’-O-ribose methylation, a pathway coronaviruses are sensitized to by inhibiting nsp16’s 2’-O-ribose methyltransferase activity,[77, 78] and DUSP11 with 5’-triphosphatase and diphosphatase activity can either directly sensitize viral RNA to exoribonucleases as with hepatitis C virus (HCV)[79] or control RIG-I responses by altering viral or host triphosphate RNA balances.[80, 81] RNA helicases are relatedly enriched yet as with 3CLpro cleavages have varying proviral and antiviral effects. Novel PLpro cleavages include neuron navigator 2, which possesses both helicase and exoribonuclease activity and is expressed in sensory nervous tissue susceptible to coronavirus infection, and DEAD box protein DDX42, which was recently shown to have broad antiviral activity including SARS-CoV-2 inhibition.[82]

Lastly, as with 3CLpro,[1] endogenous retroviral proteins were enriched in cleavages, yet HERV-K genes were recently found to be more expressed in tracheal aspirates of patients with severe COVID-19 compared to uninfected and mildly infected patients.[83] Because this expression correlates with mortality, it is worth investigating inhibiting this mechanism with antiretroviral drugs commonly used against HIV.[84] HIV protease inhibitors cannot directly inhibit coronavirus proteases at reasonable concentrations due to their difference in mechanism,[85] and lopinavir-ritonavir did not improve outcomes of patients hospitalized with COVID-19,[86] indicating that targeting HERVs in COVID-19 may be difficult or too indirect for therapeutic efficacy.

## Conclusion

Many results from my previous work on 3CLpro cleavages including enrichment of epithelium and brain, depletion of olfactory receptors, and enrichment of ubiquitination, viral RNA stabilizing and degrading mechanisms, and endogenous viral proteins are preserved in PLpro predictions.[1] Combination of the two cleavage datasets allowed for additional enrichment analysis of known substrates of cleaved E3 ubiquitin ligases, revealing that coronaviruses can regulate ubiquitination in host pathways independent of direct PLpro deubiquitinating activity. Experimental verification of these cleavages and characterization of the time dependence of these ubiquitination effects may reveal novel drug targets against COVID-19 or future human coronaviruses. Lastly, expansion of the training dataset to more divergent PLpro-containing Nidoviruses may be helpful in developing methods to phylogenetically weight aligned cleavage datasets to maximize accuracy for viral lineages or hosts of interest.

## Acknowledgements

I am very grateful for my mother, Victoria Prescott, Esq., and friends who have given me invaluable help and advice throughout my work on this project.

